# Profiling the intestinal microbiota, plasma bile acids and inflammation markers reveals novel associations in Crohn’s disease and Ulcerative colitis

**DOI:** 10.1101/2024.04.08.588526

**Authors:** Stefanie Prast-Nielsen, Anna Löf Granström, Ali Kiasat, Gustav Ahlström, Gabriella Edfeldt, Susanne Rautiainen, Fredrik Boulund, Fredrik O Andersson, Johan Lindberg, Ina Schuppe-Koistinen, Ulf O Gustafsson, Lars Engstrand

## Abstract

**Background and aims:** Our study explores signatures for Crohn’s disease (CD) and Ulcerative Colitis (UC) reflecting the interplay between the intestinal microbiota, systemic inflammation, and plasma bile acid homeostasis.

**Methods:** 1,257 individuals scheduled for colonoscopy were included and completed a comprehensive questionnaire. Individuals with IBD (‘CD’ n=64 and ‘UC’ n= 55), were age– and gender-matched to controls without findings during colonoscopy. Taxonomic profiles of the fecal microbiota and plasma profiles of inflammatory proteins and bile acids were used to build disease classifiers. Omics integration identified associations across datasets.

**Results:** *B. hydrogenotrophica* was associated with CD and *C. eutactus, C. sp. CAG 167, B. cellulosilyticus, C. mitsuokai* with controls. Ten inflammation markers were increased in CD, and eleven bile acids and derivatives were decreased in CD, while 7a-Hydroxy-3-oxo-4-cholestenoate (7-HOCA) and chenodeoxycholic acid (CDCA) were increased compared to controls.

In UC, commensals such as *F. prausnitzii* and *A. muciniphila* were depleted. CCL11, IL-17A, and TNF were increased in UC and associated to gut microbial changes. Correlations between taxa and bile acids were all positive.

**Conclusions:** For both CD and UC, taxonomic differences were primarily characterized by a reduction in commensal gut microbes which exhibited positive correlations with secondary bile acids and negative correlations with inflammation markers, potentially reflecting protective mechanisms of these commensal microbes.

## Introduction

Inflammatory Bowel Disease (IBD) is a persistent immune-mediated condition affecting the intestines, characterized by two primary types: Ulcerative colitis (UC) and Crohn’s disease (CD). As a multifactorial disease, the etiology and pathogenesis of IBD are complex and remain unestablished. The gut microbiota plays a crucial role in sustaining homeostasis within the intestines, and both compositional and functional disruption to this microbial community is recognized as a significant contributor to IBD^1,2^. The host and gut microbiota interact and produce a wide range of metabolites or compounds either from anaerobic fermentation of undigested dietary components, such as short-chain fatty acids (SCFAs), or endogenous substances generated by the host, such as bile acid metabolites^3,4^. Primary bile acids are synthesized from cholesterol in the liver and then transported to the intestine, where they are converted into secondary bile acids by specific bacterial species (e.g., *Clostridium* and *Eubacterium*)^3^. In UC patients, gut dysbiosis has been found to induce secondary bile acid deficiency, promoting a pro-inflammatory state^5^.

Primary bile acids have been found elevated in fecal samples from IBD patients, while secondary bile acids were decreased^6,7^. The gut microbiota plays a pivotal role in bile acid metabolism; however, it remains unclear which microbes influence plasma levels of which specific bile acids in IBD patients. A disruption in the microbial homeostasis may also lead to local inflammation and a disruption of the protective intestinal barrier leading to the phenomenon called “leaky gut”. This may cause upregulation of systemic inflammation proteins years prior to an IBD diagnosis^8^.

We here aimed to identify specific signatures for CD and UC in the intestinal microbiota, plasma bile acids and inflammation profiles and investigated associations between gut microbes and systemic aberrations in patients with CD and UC.

## Material and methods

### Study design and Setting

The current cross-sectional study invited all individuals referred for colonoscopy at Danderyd Hospital (a tertiary center in Stockholm) during the study period, November 1^st^, 2016, and July 1^st^, 2019, for participation. Stool samples were collected before bowel preparation with Movprep®. Plasma samples were collected for analysis of bile acids and inflammatory proteins the same day as the colonoscopy.

### Participants

The study flowchart and number of participants in each step is shown in Figure 1. Of the 2,395 individuals asked to participate in the study, 1,259 (52.6%) agreed on participating. Two individuals were excluded due to interrupted colonoscopy. All individuals included (n = 1,257) gave written informed consent before participating.

**Figure 1.**
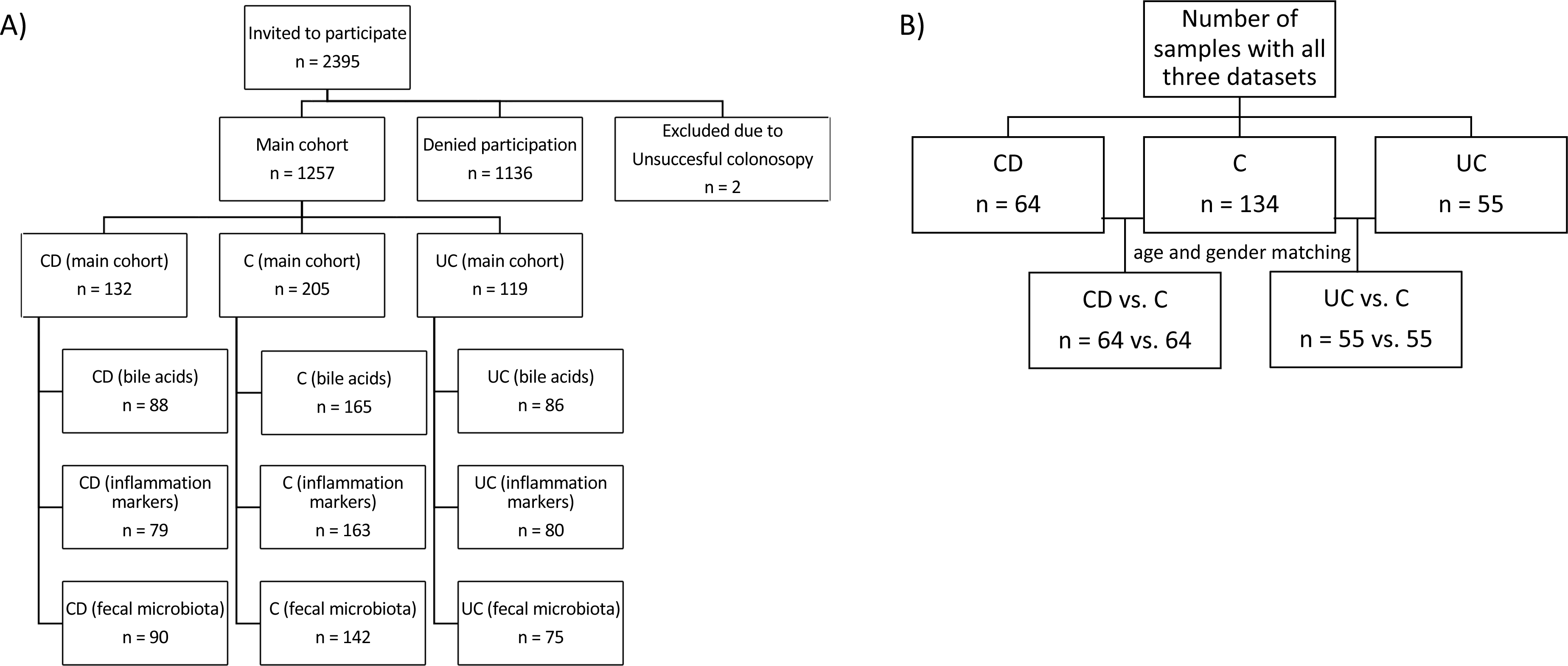
Consort diagram of the study population: A) The main cohort consisted of 1257 individuals including 132 individuals with CD, 205 individuals with a clean colon (C), and 119 individuals with UC. The remaining individuals had other gastrointestinal diagnoses. Bile acids and inflammation markers were measured in plasma of included individuals with no antibiotics up to three months prior to sampling. Fecal samples were shotgun sequenced for microbial profiles. B) For the analyses in this study, we included only individuals with all three –omics datasets and matched both CD and UC patients to C by age and gender. C = control. CD = Crohn’s disease. UC = Ulcerative colitis.

Within the scope for the current study, individuals with IBD (‘CD’ n=64 and ‘UC’ n= 55), were age– and gender-matched to control individuals with no findings on colonoscopy and no history of colorectal cancer or polyps (C) n=64 or n=55, respectively).

### Exposures

Information on IBD diagnosis, ‘CD’ or ‘UC’, and severity of disease was assessed by the examining endoscopist, and graded in accordance with Simplified endoscopic activity score point system for CD (SES-CD) and Mayo endoscopic sub score for UC^9,10^. For subgroup analysis, IBD groups were further divided into ‘CD remission’ (SES-CD 0-2) and ‘CD active disease’ (mild (SES-CD 3-6), moderate (SES-CD 5-15), or severe disease (SES-CD>15) and ‘UC remission’ (Mayo 0) or ‘UC active disease’ (mild (Mayo 1), moderate (Mayo 2) and severe disease (Mayo 3).

### Outcomes

#### Metagenomic sequencing

Stool samples were thawed and vortexed, 800 µl were transferred to a ZR BashingBead lysis tube (0,1 & 0,5 mm). A positive control of 75 µl of ZymoBIOMICS Community standard in 725 µl DNA/RNA shield and a negative control of 800 µl of DNA/RNA shield were included. All samples were bead beat in a FastPrep24 5G, 5 times at 6,0 m/sec for 1 min with 5 min of pause between beatings. The samples were centrifuged at 10,000 g for 1 min and 200 µl of the sample was added to a deep well plate for purification in a Tecan Fluent according to ZymoBIOMICS 96 MagBead DNA protocol followed by elution with 75 µl EB buffer (Qiagen). The eluted DNA was stored at –20° C until further analysis.

Fifty ng of genomic DNA was used for library preparation with MGÍs FS library prep set (MGI, Shenzhen, China) according to manufacturer’s instructions. The libraries’ quality was confirmed with the TapeStation D1000 kit (Agilent, USA) and the libraries quantity was assessed using QuantIT HighSensitivity dsDNA Assay on (Thermofisher, USA) using a Tecan Spark (Tecan, Switzerland). Circularized DNA of equimolarly pooled libraries were prepared using MGI Easy Circularization kit (MGI Tech). DNBseq 100 bp paired-end sequencing was performed using a DNBSEQ-T7 sequencing instrument (MGI, Shenzhen, China) according to the manufacturer’s instructions.

#### Sequencing quality control and taxonomic profiling of metagenomic data

Sequencing data was processed using StaG-mwc, a Snakemake-workflow for reproducible metagenomic sequencing analysis, version 0.4.3.^11^ Within this workflow, fastp version 0.20.0 was used for quality control and filtering. Kraken2 version 2.0.8_beta for removal of human reads aligning to the human genome version GRCh38. Only samples with more than 20 million non-human reads of high quality were used for further analysis. Taxonomic profiling was performed using MetaPhlAn version 3.0.7 with default settings including “unclassified” estimation and spurious taxa were filtered.

#### Bile acid profiling

Cold methanol containing an internal standard (IS) mix of ten deuterated bile acids was added to thawed plasma samples to precipitate the proteins. Samples were vortexed and centrifuged (10 min). The supernatant was evaporated to dryness and reconstituted with 10% acetonitrile in ammonium acetate solution. Quality control samples (QC) of human plasma containing all measured substances were prepared using the same procedure. Calibration mixture containing 20 bile acids was diluted to six concentrations in the range of 50-5000 nM.

Extracted plasma samples were separated on an Acquity HSS T3 column using a Waters Acquity UPLC (Ultra Performance Liquid Chromatography) system with gradient elution using ammonium acetate solution and acetonitrile. Mass spectrometry (MS) data was collected using Waters Xevo TQS™ mass spectrometers (triple quadrupole), electrospray ionization (ESI) in negative ion mode and multiple reaction monitoring (MRM) for 25 known and 38 tentative bile acid was used. Target Lynx (Waters) was used for peak integration. The response (sample peak area/IS peak area) for each bile acid in the sample was normalized against the corresponding response in the QC sample.

In a follow-up to the results from the machine learning models, an additional analytical round was performed on a selected number of bile acids, which were shown to be importan features in the models. Twelve known bile acid, of which eight with calibrator and four tentative bile acid were analysed. Some of the tentative bile acids from the first analytical round were identified before the follow-up round.

#### Profiling of inflammatory proteins

Plasma proteins were measured using the Olink® INFLAMMATION panel (Olink Proteomics AB, Uppsala, Sweden) according to the manufacturer’s instructions. The Proximity Extension Assay (PEA) technology enables 92 proteins to be analyzed simultaneously, using 1 µL of sample.^11^ Data was quality checked and normalized using an internal extension control and an inter-plate control, to adjust for intra– and inter-run variation. The final assay read-out is presented in Normalized Protein eXpression (NPX) values, which is an arbitrary unit on a log2-scale where a high value corresponds to a higher protein expression. All assay validation data (detection limits, intra– and inter-assay precision data, etc.) are available on the manufacturer’s website (www.olink.com).

### Covariates

#### The lifestyle questionnaire

Study participants completed a validated 13-page questionnaire with 277 questions to collect information on dietary-, lifestyle– and bowel habits (including Bristol Stool Scale (BSS)).^12^ Data on gender, age, body mass index (BMI), smoking (smoker/nonsmoker) and education level were included here. Out of 456 individuals included in the study, 451 (98,9%), completed the questionnaire. Educational level was categorized into ≤9 years, 10-12 years and ≥13 years of schooling. Bristol Stool Scale was categorized into the following groups: slow (BSS 1-4), normal (BSS 3-4 only), rapid (BSS 3-7) and varied (BSS both 1-2 and 5-7). Psychiatric scores (PHQ-9 for depression, GAD-7 for anxiety, PSS-4 for stress) were also collected. Individuals using antibiotics less than three months before colonoscopy were excluded.

#### Dietary components

From the lifestyle questionnaire, data on dietary intakes were collected. All nutrients were energy-adjusted to the mean energy intake in the study population using the residual methods.^13,14^ The Alternate Healthy Eating Index (AHEI) was calculated from the questionnaire.^15^ The AHEI includes 10 components, where each of the following items were given a score between 0 and 10 proportionally to dietary intake: 1) vegetables, 2) fruit, 3) whole grains, 4) nuts and legumes, 5) saturated fatty acids, 6) polyunsaturated fatty acids), 7) alcohol consumption, 8) sugar sweetened drinks and fruit juices, 9) red and processed meat, and 10) sodium. A higher score represents a higher adherence to the AHEI.

#### Bioinformatics/statistical analyses and machine learning

All statistical analyses and visualizations were performed in R version 4.2.2. Matching for gender and age was performed using the MatchIt library version 4.5.0. We used centered log ratio (clr)-transformed relative abundance values for statistical analysis of the metagenomics dataset. For Permutational Multivariate Analysis of Variance (PERMANOVA), the adonis function of the vegan package version 2.6-4 was used with euclidean distances and 99,999 permutations. Samples with missing data from the questionnaire were excluded here. FDR (false-discovery rate)-values were calculated from p-values and covariates with an FDR < 0.1 were considered significant.

The following machine learning algorithms, glmnet, pcaNNet, Ranger Random Forest, knn, XgbTree, and svmRadial were run and optimized using the caret package version 6.0-93 in R. For omics integration of taxa and bile acid profiles or inflammation markers, respectively, PLS-DA was performed using DIABLO (Data Integration Analysis for Biomarker discovery using Latent variable approaches for Omics studies) within the mixOmics R package.^17,18^ Further details on the bioinformatics analyses can be found in the supplementary information.

### Ethical Approval

The research protocol was approved 2016-02-03 by the Karolinska Institute Ethics Committee (2015/2138-31/2) and carried out in accordance with the declaration of Helsinki of the World Association (1989) (clinical trial number: NCT03302715, www.clinicaltrials.gov).

### Data availability

All data files used as input for the analyses are provided as supplementary files. Sequencing data will be published in the European Nucleotide Archive as soon as the article has been accepted for publication.

## Results

### Study population characteristics

For individuals with CD vs. C, samples were age-matched (n_CD_=64, n_C_=64,) and for UC vs. C, samples were both age– and gender matched (n_UC_=55, n_C_=55,) as these variables were significantly different between cases and controls before matching.

Basic characteristics are shown in Table 1. Laboratory samples, educational level and dietary intake were similar between groups while the proportion of previous surgery was highest among individuals with CD (18.7%). Among CD patients, 65.6% were in clinical remission vs 90.9% in UC, which was reflected in ongoing medication with immunomodulators and biological treatment, 31.3% and 37.5% vs 18.2% and 10.9% respectively.

### Sequencing results

Mean sequencing depths in each group was as follows: For the CD cohort: C 109 Mreads; CD: 113 Mreads and for the UC cohort: C 101 Mreads; UC: 101 Mreads.

### Identification of factors significantly influencing the overall profiles of each dataset

Permutational Multivariate Analysis of Variance (PERMANOVA) for covariates is summarized in Table 2 for both cohorts and all datasets. For the CD/C cohort, several disease-associated covariates were significant for the bile acid profiles and inflammation profiles but not the taxonomic profiles, whereas in the UC/C cohort; the IBD diagnosis was only significant for the taxonomic profiles. Age, gender, and BMI were significant in the PERMANOVA analysis for various profiles in both cohorts but not significantly different between groups after matching. Medications influenced overall diversity in datasets of both cohorts.

### Determining the optimal machine learning algorithm to discriminate individuals afflicted with Crohn’s disease vs controls

Several machine learning algorithms were applied to analyze which algorithm could best predict disease in a separate test set of our cohort which was not used for training the models. As input, each dataset, i.e., taxonomic (taxa), bile acids (bile) and inflammatory protein profiles (proteins) were used separately, as well as combined (all). The results are summarized in Table 3 with highest performance marked in bold. The features contributing to the best performing model were ranked by importance and assigned a negative value if they were associated with a control and a positive value if they were associated with CD (Figure 2). Four out of five species were associated with C, one with CD. All ten inflammation markers were associated with CD, and eleven bile acids and related metabolites were decreased in CD, while three were increased. The best model performances were achieved with either bile acid profiles alone or a combination of all datasets. Using a combination of all datasets, five features were sufficient to achieve an accuracy of 0.82 in predicting disease in our test set. These features were identical to the top two species in the taxonomic model, i.e., *Coprococcus eutactus* and *Clostridium sp. CAG 167* associated with controls, and the top three CD-associated inflammation markers TNF, CCL20 and CCL11.

**Figure 2.**
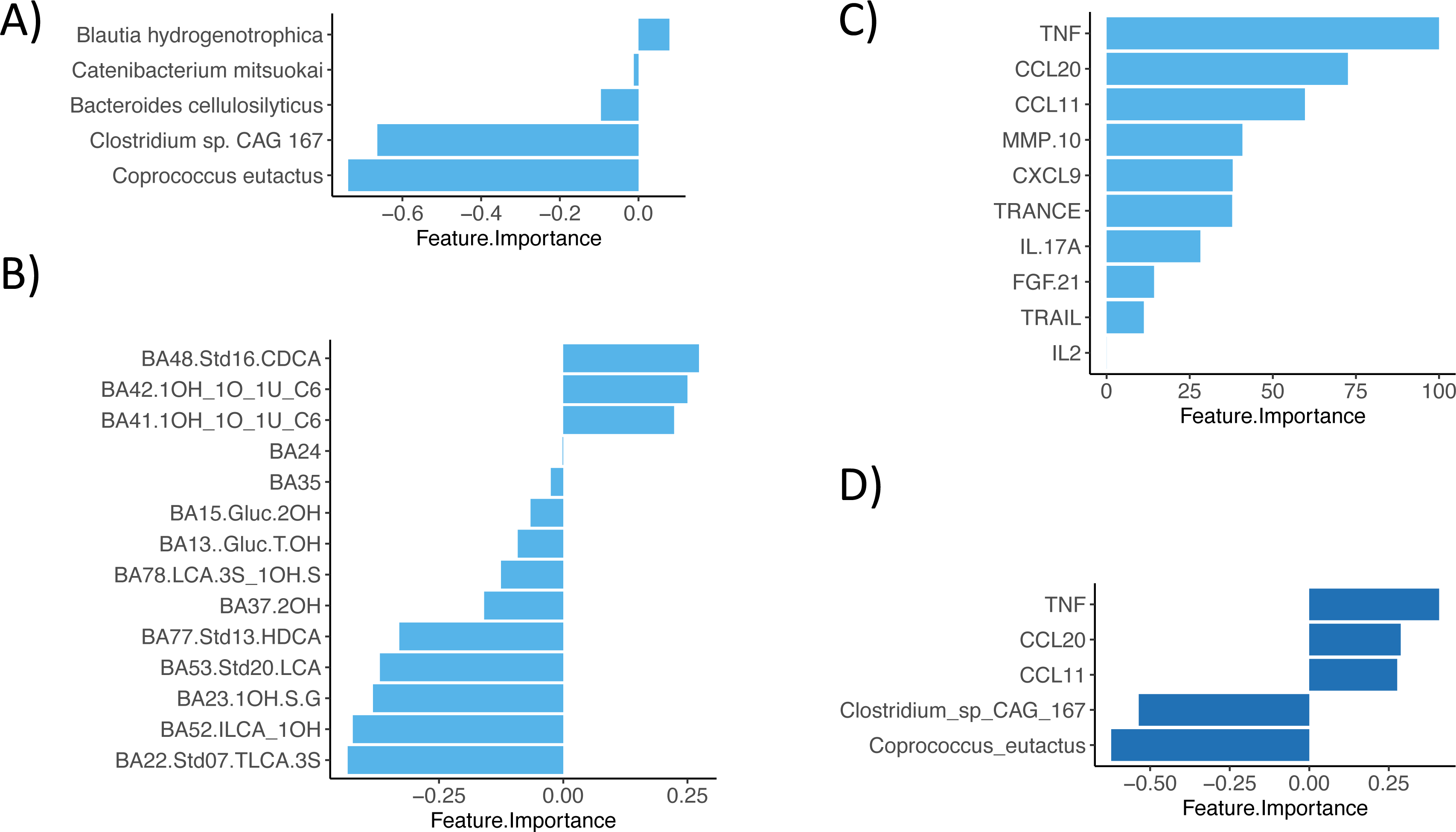
Discriminant features (C vs. CD) of the best performing model for each dataset (A-C) and all combined datasets (D). Features ranked by importance with positive values associated with CD and negative values associated with C for A) taxonomic data, B) bile acids, C) inflammation proteins, and D) all datasets combined.

### Integrative –omics analysis reveals associations between gut microbiota and plasma bile acids and inflammation markers in Crohn’s Disease

Using DIABLO, we analyzed which intestinal species might drive changes in plasma bile acid profiles (Figure 3 A) and inflammation (Figure 3B). Full datasets were used for the integrations without feature selection based on our prior analysis to enable full analysis of all possible associations. The model for the microbiota and systemic inflammation achieved a performance accuracy of 0.71 and revealed positive and negative associations. Six bacterial species associated with controls were negatively associated with CCL11 and/or CCL20 – top inflammation markers in the models above, whereas *Flavonofractor plauti* and *Ruminococcus gnavus* showed positive correlations with these. Positive associations dominated the integration model of microbes with bile acids which achieved an accuracy of 0.64. Seven species correlated positively with six bile acids with a correlation coefficient above 0.4. Of these, five species, i.e., *Ruminococcaceae bacterium D5, Clostridium sp CAG 167, Roseburia sp CAG 309, Firmicutes bacterium CAG 95* and *CAG 238* were negatively associated with inflammation (Figure 3A). In contrast, both species positively associated with inflammation showed negative correlations to these bile acids, which included secondary bile acids and derivatives such as iso-lithocholic acid (ILCA), taurolithocholic acid-3-sulfate (TLCA-3S) and hyodeoxycholic acid (HDCA) (Figure 3B).

**Figure 3.**
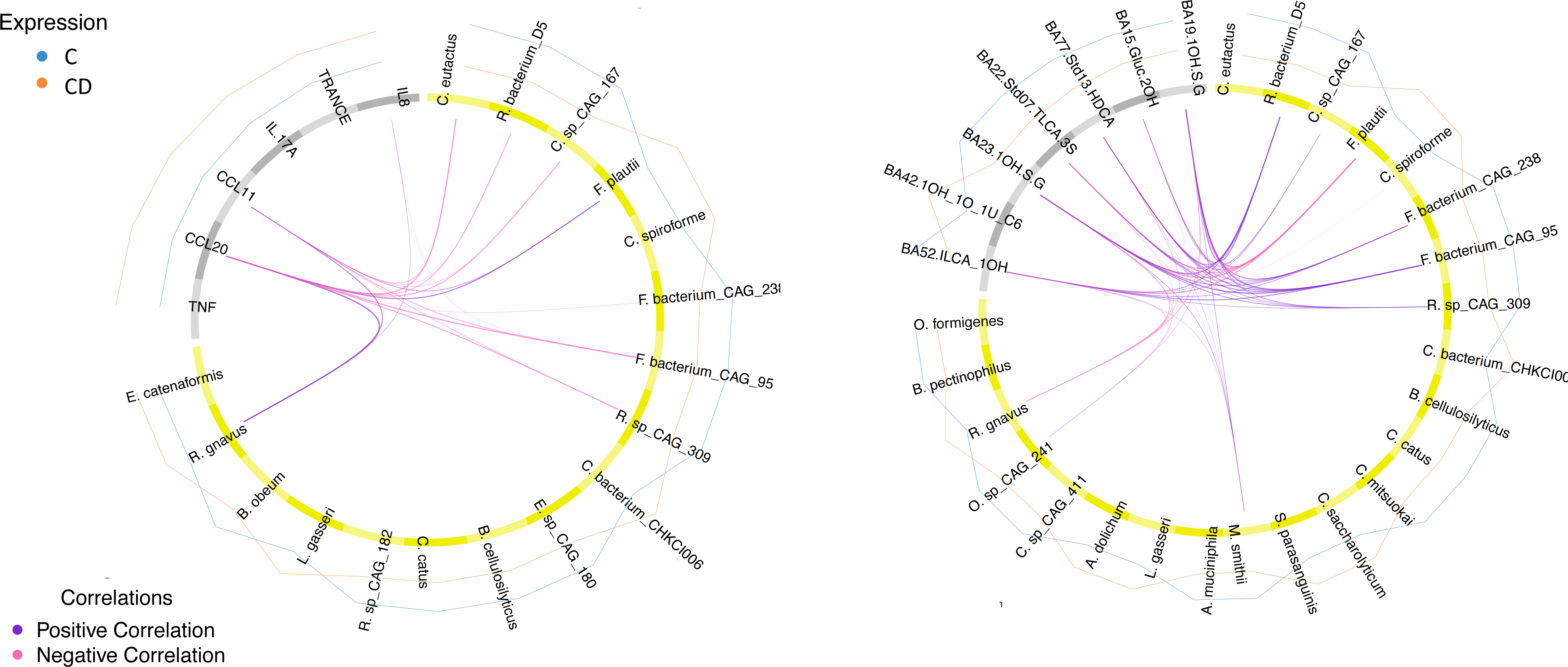
Circos plots of DIABLO results for A) integration of taxonomic and inflammation profiles and B) integration of taxonomic and bile acid profiles in C vs CD. Correlation coefficients above 0.4 are displayed in purple for positive and in pink for negative associations across datasets. The bacterial species are shown in yellow and the inflammation proteins and bile acids in gray. Group levels of each feature are represented by blue lines for C and orange lines for CD. Model accuracy for A) was 0.71, for B) 0.64.

### Determining the optimal machine learning algorithm to discriminate individuals afflicted with Ulcerative Colitis vs controls

The same machine learning algorithms used for the CD sub-cohort were used for the UC sub-cohort for each dataset, i.e., taxa, bile, inflammatory proteins, and a combination of all. The resulting accuracies for classification of the independent test set are shown in Table 4. The best model was achieved for the inflammatory proteins with 0.78 in accuracy using 20 features, followed by the metagenomic data with 0.72 using 10 features. Of these selected ten taxa, the majority were associated with controls. Neither of the algorithms tested achieved a good model for the bile acid profiles, the highest being PLS-DA with 0.63 in accuracy using five features (Figure 4B).

**Figure 4.**
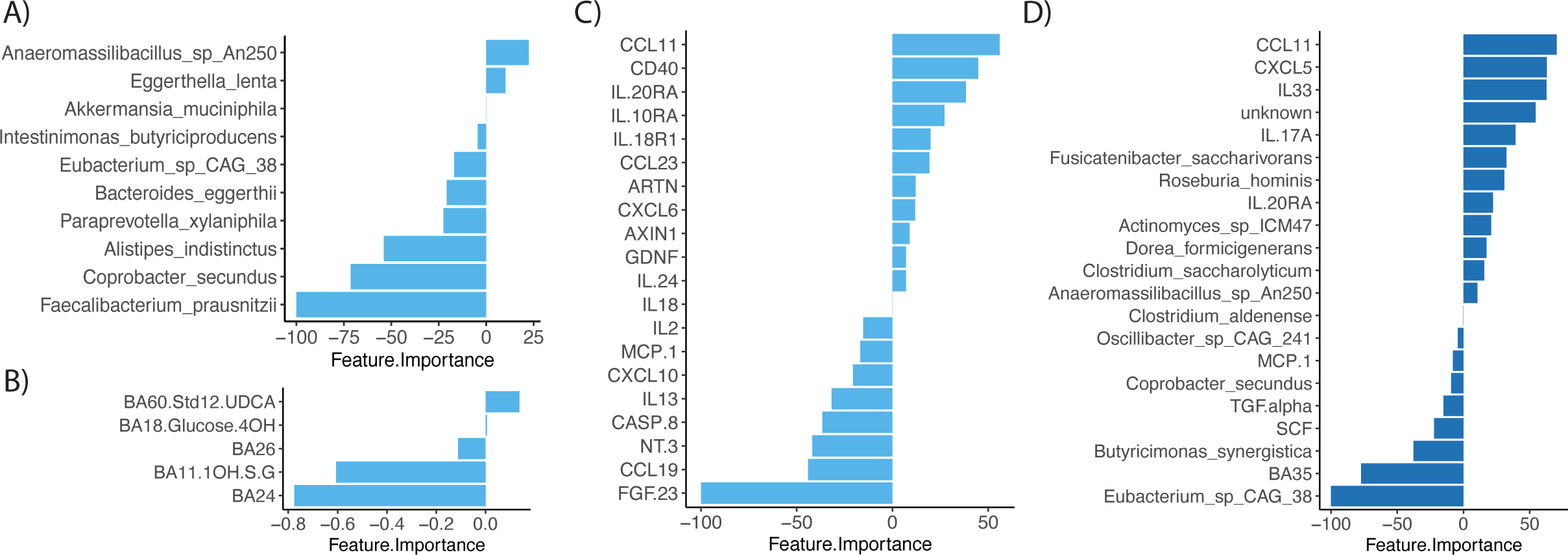
Discriminant features (C vs. UC) of the best performing model for each dataset (A-C) and all combined datasets (D). Features ranked by importance with positive values associated with UC and negative values associated with C for A) taxonomic data, B) bile acids, C) inflammation proteins, and D) all datasets combined.

Twelve out of twenty important inflammation markers in our final model were higher in UC patients, including typical UC-associated peptides such as CCL11, CD40, IL-20RA, IL10RA and IL18R1 (Figure 4C). Eight proteins were decreased in our UC patients. For the combined dataset of taxonomic, bile and inflammation profiles, the accuracy of the final model was 80.1 using 21 features. Of these, only one was a bile acid of unknown structure (BA35) (Figure 4D). Eight features were inflammation-associated proteins which also were amongst the important features of the model built only using inflammatory proteins. Most features in this model were gut microbial species (12/21). Several were associated with UC, however, the overall most important feature *Eubacterium sp. CAG 38*, was associated with controls.

### Integrative –omics analysis reveals associations between gut microbiota, plasma bile acids and inflammation markers in Ulcerative Colitis

DIABLO was used to analyze which intestinal species might drive changes in bile acid profiles (Figure 5A) and inflammation (Figure 5B) in the UC sub-cohort. All features in each dataset were included to allow full analysis of all possible associations. Performance of the model integrating taxonomic and inflammation data, reached an accuracy of 0.65. The majority of correlations between taxa and inflammation markers were negative. The only positive correlation detected above 0.4 was between *C. spiroforme* and IL-17A. Gut microbial species which were negatively associated with CCL11, IL-17A and TNF were all depleted in UC patients. Of these, *Eubacterium sp CAG 38* and *Akkermansia muciniphila* were the most discriminative features. These two taxa were also the most discriminative and associated to various, predominantly secondary, bile acids in an integration model with species and bile acid profiles. This model reached an accuracy of 0.60 and all correlations were positive (Figure 5B).

**Figure 5.**
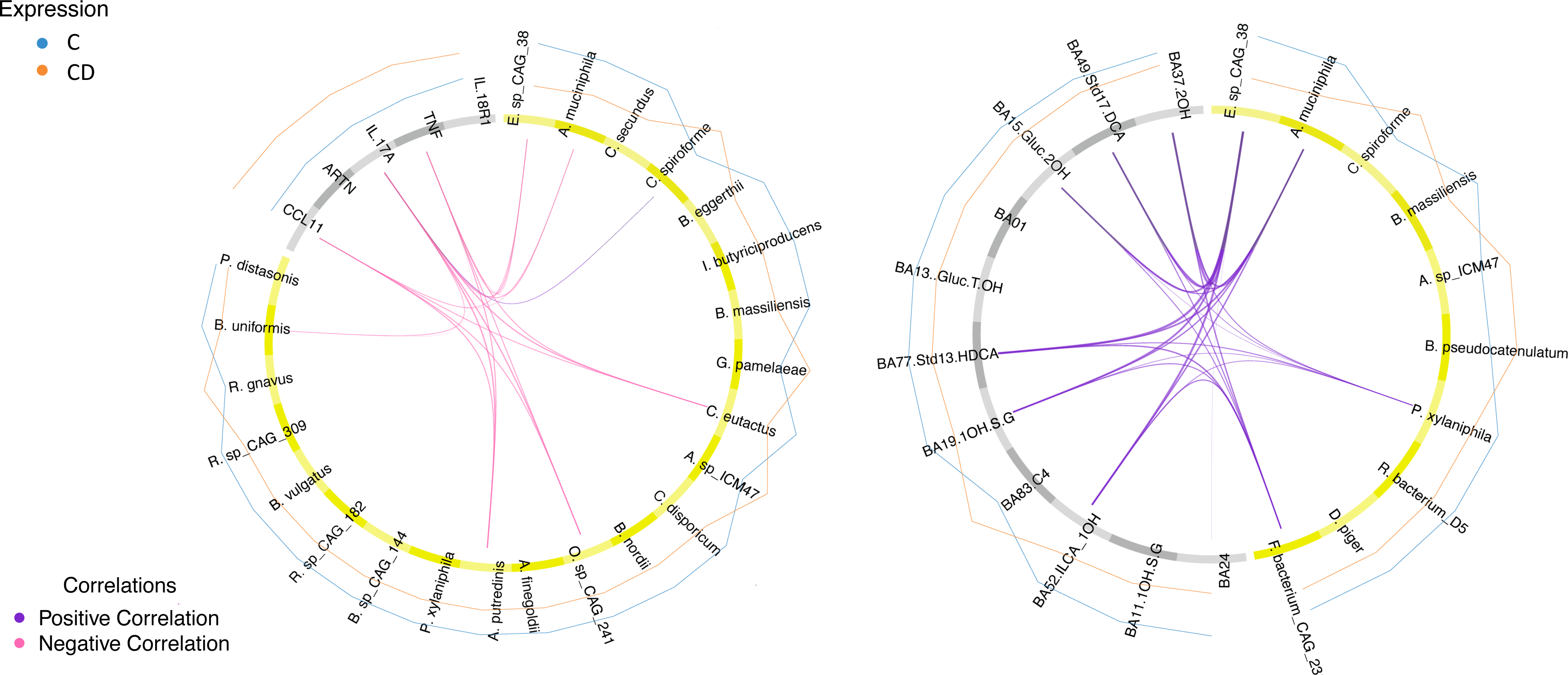
Circos plots of DIABLO results for A) integration of taxonomic and inflammation profiles and B) integration of taxonomic and bile acid profiles in C vs UC. Correlation coefficients above 0.4 are displayed in purple for positive and in pink for negative associations across datasets. The bacterial species are shown in yellow and the inflammation proteins and bile acids in gray. Group levels of each feature are represented by blue lines for C and orange lines for UC. Model accuracy for A) was 0.65 and for B) 0.6.

## Discussion

In this cross-sectional study, machine learning algorithms were utilized to analyze and characterize the intestinal microbiota, inflammation markers, and plasma bile acids in a group of 64 individuals with Crohn’s disease and 55 individuals with Ulcerative colitis. These groups were compared to appropriately matched control individuals. This comprehensive approach revealed novel associations. For CD, all inflammatory protein features were associated with disease, while different taxonomic features and bile acids were associated with either disease or control. The majority of commensal taxonomic features exhibited a negative association with inflammation markers while showing a positive association with bile acids. For UC individuals we identified both taxa, bile acids and inflammation markers which were associated with either disease or controls. Most correlations between taxa and inflammation markers were negative, whereas positive for all bile acids.

### Crohn’s disease

The finding that the taxonomic feature, *Coprococcus eutactus* was associated with controls rather than CD is consistent with previous research. *C. eutactus* has been shown to decrease in individuals with pro-inflammatory diets and Crohn’s disease.^20^ However, it increased when disease improved, and exhibited a negative correlation with inflammatory markers.^21^ Furthermore, the recently identified microorganism, *Clostridium sp. CAG 167*, that has been recognized as one of the most beneficial bacteria within the gastrointestinal system, was associated with controls in the current study. Inflammatory markers were positively associated with CD including the topmost important markers TNF, CCL 20 and CCL11.^21^ The cytokine TNF is an inflammatory mediator that has been targeted for medical treatment of CD, the chemokine CC-chemokine-ligand-20 (CCL-20) has previously been identified as a promotor of colitis associated colorectal cancer (5) and CCL11 induced exacerbation of colitis in a mouse model (Polosukhina et al. 2020).^22–24^

Eleven bile acids were negatively and three were positively associated with CD. Two of the bile acids associated with CD indicated as BA41 and BA42 were subsequently identified as 7a-Hydroxy-3-oxo-4-cholestenoate (7-HOCA), which showed a double-peak in the chromatogram. 7-HOCA has, to our knowledge, not been associated with CD previously. The remaining bile acid associated with CD was chenodeoxycholic acid (CDCA) which has previously been shown to be elevated in CD.^25^ Both 7-HOCA and CDCA are primary bile acids and did not correlate with the microbiota in our analysis. Bile acids associated with controls included the secondary bile acid lithocholic acid (LCA), a product of CDCA, and its derivatives. Of these, TLCA-3S, ILCA and glycolithocholic acid 3-sulfate (GLCA-3S) were not only the most important features in the bile acid classification model but also showed strongest positive correlations to members of the gut microbiota.

All species positively correlating with bile acids were also negatively associated with inflammation, thus, a depletion of these bacteria may disturb bile acid metabolism and promote inflammation. Restoration of these species might alleviate these disturbances and contribute to disease improvement.

### Ulcerative colitis

Among the inflammatory markers associated with UC in this cohort, the chemokine CCL11, macrophage-related protein CD40, the cytokines IL-20RA, IL10RA and IL18R1, have all previously been described as inflammation proteins involved in the pathogenesis of UC.^26–29^ Gut microbial species which were negatively associated with CCL11, IL-17A and TNF were all depleted in UC patients in the current study. This included *Eubacterium sp. CAG 38* which had not previously been linked to UC to our knowledge. However, the genus Eubacterium has been found in reduced abundances among individuals with CD.^30^ Additionally, the mucolytic bacterium *Akkermansia muciniphila*, believed to offer protection against colitis, was found to be diminished within the UC group.^31^ Although the model integrating the bile acid profile and taxonomic data had a low accuracy of 0.60, E. *sp. CAG 38* and *A. muciniphila* showed the strongest correlations with bile acids. The majority of the bile acids that correlated positively with gut species in UC were found in the CD model as well, except DCA and 3a,12b-dihydroxycholanoic acid which showed stronger microbial associations in UC than CD.

For both CD and UC, a general pattern was observed. The taxonomic changes were dominated by the depletion of commensal gut microbes, such as *C. eutactus* and *Clostridium sp. CAG 167* for CD, and *F. prausnitzii* and *A. muciniphila* for UC. Commensal gut microbes also showed the strongest positive associations with secondary bile acids like ILCA and HDCA, and negative associations with inflammation markers, like CCL11, in both subtypes of IBD suggesting a depletion of several commensal species may lead to increased inflammation and perturbation of bile acid metabolism in patients with IBD.

### Machine learning algorithms

In the current study machine learning algorithms were used for the analysis of several –omics datasets. The benefits of machine learning include the possibilities to uncover complex and non-linear patterns in high-dimensional data that might not be apparent through conventional methods. These algorithms can predict outcomes or associations, aiding in identifying potential biomarkers or indicators of diseases. Finally, although machine learning can establish correlations, inferring causation from observational data cannot be done.

### Strengths and limitations

This study has a notable advantage in employing the UPLC technique to analyze plasma samples. UPLC offers superior chromatographic resolution compared to the HPLC (High-Performance Liquid Chromatography) method used in prior investigations of IBD and plasma bile acid levels. This heightened resolution allows for the differentiation of 63 distinct bile acids and derivatives.

We here combine several –omics analyses on samples from the same individuals allowing for improved disease classification by machine learning algorithms. Integration of these datasets provided novel hypotheses for how the depletion of commensal microbes may disturb bile acid metabolism and fuel inflammation.

In contrast to earlier studies, our analyses encompass a comprehensive questionnaire with an impressive compliance rate of 98.9%. This questionnaire captures detailed information regarding participants’ dietary choices, lifestyle, and bowel habits. Alongside this, the study benefits from other objectively collected clinically significant data, such as endoscopic disease activity status. Collectively, these factors enhance the robustness of the study.

Nevertheless, the study has limitations. This is a cross-sectional study which only allows for associations and cannot infer causation. Prior to obtaining blood samples, all participants fasted and underwent bowel preparation using Movprep® which may lead to changes in bile acid metabolism as the body adjusts to a temporary lack of incoming nutrients. Additionally, individuals in the control group were selected based on the absence of pathological findings during colonoscopy. However, given that they were referred for colonoscopy due to specific reasons, they cannot be unequivocally classified as entirely healthy. Most individuals were on medication, including IBD medications and proton pump inhibitors known to influence the composition of the gut microbiota. Almost all UC patients were in remission, while 34% of the CD patients experienced active inflammation at the time of sample collection.

### Interpretation of the Key Findings

For both CD and UC, taxonomic profiles were primarily characterized by a reduction in commensal gut microbes. These microbes exhibited the strongest positive correlations with secondary bile acids and negative correlations with inflammation markers. This suggests that the depletion of several commensal species may contribute to heightened inflammation and disruption of bile acid metabolism in patients with inflammatory bowel disease (IBD). This study also highlights potential biomarkers, especially for CD, such as the two primary bile acids, 7-HOCA and CDCA, which warrant further investigation in future studies.

This work was supported by grants provided by Ferring Pharmaceuticals and Danderyd Hospital, Stockholm County.

## Acknowledgements

We thank all participants of the study and all clinical personnel involved. We extend our thanks to Tatjana Pavlenko at Uppsala University, Professor in Statistics, with expertise on high-dimensional problems in statistical machine learning, detection, and identification of sparse signals, for advice on analysis appraoched. We acknowledge the Affinity Proteomics-Stockholm Unit at SciLifeLab for technical support and generation of protein data.

## Disclosures

S.P.-N., G.A., G.E., F.B., I.S.-K., and L.E. received funding from Ferring Pharmaceuticals. All other authors have nothing to disclose.

## Author contributions

Stefanie Prast-Nielsen: Major data analysis, Interpretation of results, Manuscript writing, reviewing and editing

Anna Löf Granström: Study design, Data collection, Interpretation of results, Manuscript writing, reviewing and editing

Gustav Ahlström: Major data analysis, Manuscript writing, reviewing and editing

Ulf O Gustafsson: Study design, Data collection, Interpretation of results, Manuscript writing, reviewing and editing

Susanne Rautiainen: Interpretation of results, Manuscript reviewing and editing.

Ina Schuppe-Koistinen: Study design, Planning analyses, Manuscript writing, reviewing and editing

Lars Engstrand: Study design, Planning analyses, Manuscript reviewing and editing, obtained funding

Gabriella Edfeldt: Inflammatory protein analysis, Interpretation of results, Manuscript reviewing and editing

Ali Kiasat: Study design, Data collection processing, Manuscript reviewing Fredrik O Andersson: Bile acid profiling, Manuscript reviewing and editing Johan Lindberg: Bile acid profiling, Manuscript reviewing and editing

Fredrik Boulund: Data collection systems development, Supporting data analysis, Manuscript reviewing and editing

