## Supplemental information for "Profiling the intestinal microbiota, plasma bile acids and inflammation markers reveals novel associations in Crohn’s disease and Ulcerative colitis"

**Supplementary information**

Methods

*Bioinformatics analyses*

Datasets were autoscaled prior to multivariate analyses. Each dataset was randomly split into a training set (70% of the samples) and a test set (30% of the samples) with identical numbers of cases and controls in each set. The following machine learning algorithms, glmnet, pcaNNet, Ranger Random Forest, knn, XgbTree, and svmRadial were run and optimized using the caret package version 6.0-93 in R. The train function of caret was applied to the training set to fit predictive models over different tuning parameters with a tune length of 18. The models were tuned using a grid search and a leave-one-out cross-validation (LOOCV) approach within the trainControl function. For partial least squares discriminant analysis (PLS-DA) we used the mixOmics R package version 6.23.4 with LOOCV on the training set.

All resulting models were validated for their accuracy of correctly predicting the outcome (C or CD/C or UC) of the test set as follows:

$$\text{Accuracy of test set} = \frac{\text{True positives} + \text{True negatives}}{\text{All samples}}$$

The best performing model was then further optimized by testing algorithm specific hyperparameters to potentially improve model performance. The features contributing to the final model were visualized ranked by importance.

For omics integration of taxa and bile acid profiles or inflammation markers, respectively, PLS-DA was performed using DIABLO (Data Integration Analysis for Biomarker discovery using Latent variable approaches for Omics studies)[27],[28] within the mixOmics R package. The type of distance and the number of components for tuning of the models was chosen based on the lowest error rate achieved when running the “perf” function. A range of variables for both taxa and bile acids or inflammation markers was tested in tune.block.splsda using LOOCV. Correlation coefficients between important variables of each dataset were calculated according to Ignacio González et al.[29] and correlations were visualized in circos plots (cut-off = 0.4).
